# IPSC-derived neuronal cultures expressing the Alzheimer’s disease associated rare TREM2 R47H variant enables the construction of an Aβ-induced gene regulatory network

**DOI:** 10.1101/652446

**Authors:** Soraia Martins, Andreas Müller-Schiffmann, Martina Bohndorf, Wasco Wruck, Kristel Sleegers, Christine Van Broeckhoven, Carsten Korth, James Adjaye

**Affiliations:** Institute for Stem Cell Research and Regenerative Medicine, Medical Faculty, Heinrich-Heine University, Düsseldorf, Germany; Department Neuropathology, Heinrich-Heine University, Düsseldorf, Germany; Neurodegenerative Brain Diseases Group, Center for Molecular Neurology, VIB, Antwerp, Belgium; Department of Biomedical Sciences, University of Antwerp, Belgium

## Abstract

Recently, genes associated with immune response and inflammation have been identified as genetic risk factors for late-onset Alzheimer’s disease (LOAD). One of them is the rare p.Arg47His (R47H) variant within triggering receptor expressed on myeloid cells 2 (TREM2), which has been shown to increase the risk for developing AD 2-3-fold. Here, we report the generation and characterization of a model of LOAD using lymphoblast-derived iPSCs from patients harbouring the R47H mutation in TREM2 (AD TREM2 iPSCs), as well as from control individuals without dementia (CON iPSCs). iPSCs efficiently differentiate into mature neuronal cultures and comparative global transcriptome analysis identified a distinct gene expression profile in AD TREM2 neuronal cultures. Furthermore, manipulation of the iPSC-derived functional neuronal cultures with an Aβ-S8C dimer highlighted metabolic pathways, phagosome and immune response as the most perturbed pathways in AD TREM2 neuronal cultures. Through the construction of an Aβ-induced gene regulatory network, we were able to identify an Aβ signature linked to protein processing in the endoplasmic reticulum (ER) which emphasised ER-stress, as a potential causal role in LOAD. Overall, this study has shown that our AD-iPSC based model can be used for in-depth studies to better understand the molecular mechanisms underlying the etiology of LOAD and provides new opportunities for screening of potential therapeutic targets.

## Introduction

Currently, there are 47 million people living with dementia worldwide, a number that is estimated to increase to more than 131 million in 2050 (Patterson, 2018). Alzheimer’s disease (AD) is a neurodegenerative disease and the most common and devastating cause of dementia, contributing to 60-70% of all cases (Prince and Comas-Herrera, 2017). AD is clinically characterized by a progressive decline of cognitive functions and, according to the classical amyloid hypothesis two key molecules have been implicated in AD neuropathology: amyloid-beta (Aß) and the protein tau (Hardy and Higgins, 1992). Aß peptides are derived from sequential proteolytic cleavages of Amyloid Precursor Protein (APP). They form extracellular aggregated deposits known as amyloid plaques. Intracellularly, hyper-phosphorylated tau forms aggregates composed of twisted filaments known as neurofibrillary tangles (NFTs). As a consequence of the imbalanced crosstalk between Aß and tau, multiple neuropathological mechanism ensue, such as, synaptic toxicity, mitochondrial dysregulation and microglia-derived inflammatory responses, finally leading to neuronal death (Hardy and Selkoe, 2002; Polanco et al., 2018). Age is the greatest risk factor for AD and it can divided it into early-onset AD (EOAD) when the patients are younger than 65, and late-onset AD (LOAD) when the patients manifest symptoms after the of age 65 (Cacace et al., 2016). Despite EOAD being responsible for a small minority of all cases, the studies of familiar AD patients (fAD) have revealed important aspects of the genetic factors implicated in the disease, such as the causal mutations in *APP*, *PSEN1* and *PSEN2.* On the other hand, LOAD is a very complex and multifactorial disease where most cases are sporadic with no clear familiar pattern of disease (Bettens et al., 2013; Carmona et al., 2018). Many genetic risk factors have been implicated in increasing the susceptibility for LOAD, among which is the well establish apolipoprotein E (*APOE).* Individuals carrying one ε4 allele have a 3-fold increased risk of AD while individuals carrying the two ε4 alleles face an approximately 12-fold increased risk of AD (Bickeböller et al., 1997; Farrer et al., 1997). More recently genome-wide association studies (GWAS) have led to the discovery of other genetic variants that influence the risk for LOAD (Harold et al., 2009; Hollingworth et al., 2011; Lambert et al., 2009; Naj et al., 2011; Seshadri et al., 2010). Some of these variants are located in or near genes known to be involved in biological pathways such as cholesterol metabolism, immune response, and endocytosis/vesicle-mediated transport (Carmona et al., 2018). As a more direct link between immune responses and AD, especially microglia-related genes with an increased risk for developing LOAD were identified by high-throughput sequencing technologies (Guerreiro et al., 2013; Jonsson et al., 2013). One of multiple genetic risk variants identified in these studies is the rare p.Arg47His (R47H) variant within triggering receptor expressed on myeloid cells 2 (*TREM2*), which has been shown to increase the risk of developing AD by 2-3-fold in several European and North American populations (Cuyvers et al., 2014; Guerreiro et al., 2013; Jonsson et al., 2013; Ruiz et al., 2014; Sims et al., 2017; Song et al., 2017).

TREM2 is a cell surface receptor of the immunoglobulin superfamily expressed on various cells of the myeloid linage including CNS microglia, bone osteoclasts, alveolar and peritoneal macrophages (Colonna and Wang, 2016). Although *in vivo* ligands are currently unknown, TREM2 binds *in vitro* to anionic lipids, APOE, high-density and low-density lipoprotein, apoptotic cells and Aβ. Upon ligand binding, TREM2 initiates a signalling cascade through the association with DAP12, modulating cell proliferation and differentiation, survival, chemotaxis and inflammation (Atagi et al., 2015; Song et al., 2017; Wang et al., 2015). Importantly, TREM2 is required for microglial phagocytosis of a variety of substrates, including apoptotic neurons and Aβ and thus playing a prominent role in driving microgliosis (Colonna and Wang, 2016; Zhao et al., 2018; Zhou et al., 2018). According to neuropathology studies in AD patients, animal models, and *in vitro* studies, the *TREM2* R47H variant induces a partial loss of function of TREM2, compromising microglia function and thus contributing to the development of AD. TREM2 deficiency in AD mouse models and patients carrying the R47H variant showed decreased clustering of microglia around the plaques, thereby facilitating the build-up of Aβ plaques and injury to adjacent neurons (Jay et al., 2015; Song et al., 2018; Wang et al., 2015; Yuan et al., 2016). Recent data have shown that cells expressing the R47H variant displayed impaired TREM2-Aβ binding and altered TREM2 intracellular distribution and degradation, thus providing a potential mechanism by which *TREM2* R47H mutation increases the risk for LOAD (Lee et al., 2018; Zhao et al., 2018). Although the role of TREM2 in AD has been a focus of study using post-mortem brains, mouse models and heterologous cell lines, a consensus has not been reached probably due to the inability to effectively recapitulate this complex disease with the current models. The adoption of induced pluripotent stem cells (iPSCs) technology provides a platform to derive a reliable human disease model for better understanding the effect of risk factors in neurons derived from primary cells of affected patients. iPSC modelling of AD has provided an important proof-of-principle regarding the utility of such cells for a better understanding of the molecular mechanisms associated with the etiology of AD. So far, a number of the human iPSC-based AD models have concentrated on using iPSCs derived from EOAD or LOAD patients with unidentified mutations (Duan et al., 2014; Flamier et al., 2018; Hossini et al., 2015; Israel et al., 2012; Muratore et al., 2014; Ochalek et al., 2017; Yagi et al., 2011).

Here, we report for the first time the generation and characterization of a model of LOAD using lymphoblast-derived iPSCs from patients harbouring the R47H mutation in *TREM2*, as well as from control individuals without dementia. To date gene regulatory networks governing LOAD have been generated using human AD brain biopsies. In our current study we have shown the feasibility of using an iPSC-based approach to derive biologically meaningful pathways and an Aβ-induced regulatory network that mirrors some of the pathways that have been identified by the LOAD brain biopsies, namely immune response, phagocytosis and unfolded protein response pathways (Zhang et al., 2013). Our study thus demonstrates that AD TREM2 iPSC-derived neuronal cultures can be used for in depth studies to understand the molecular mechanisms underlying the onset of Alzheimer disease and for screening of potential therapeutic targets.

## Results

### iPSCs efficiently differentiate into a functional neuronal culture

iPSCs derived from lymphoblasts from two LOAD patients carrying the *TREM2* R47H risk variant (AD TREM2-2 and AD TREM2-4), as well as aged-matched control individuals without dementia (CON8 and CON9) were used for this study (Martins et al., 2018; Schröter et al., 2016a; Schröter et al., 2016b; Schröter et al., 2016c). The summary of the characteristics of the iPSC lines used in this study as well as their APOE status are shown in Table 1.

**Table 1.**
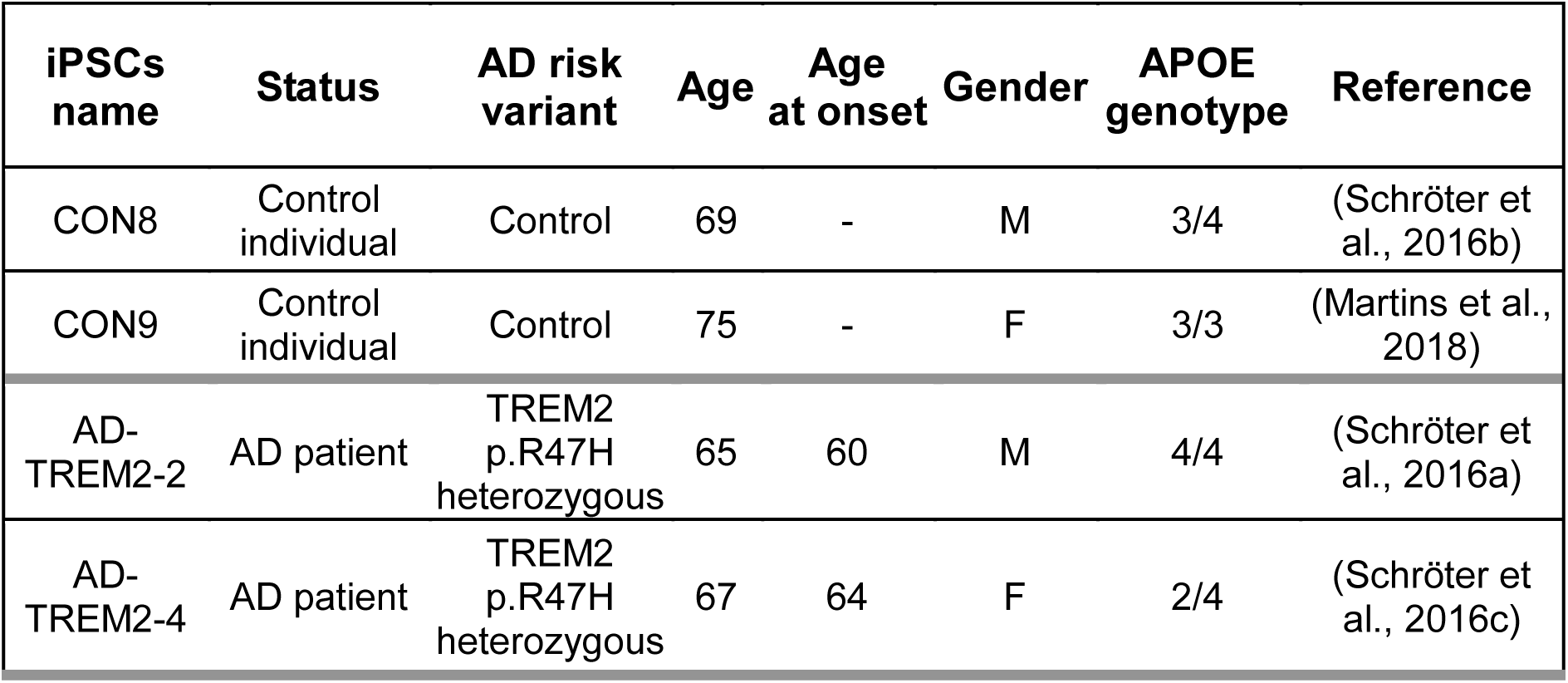
Summary of the healthy Controls and AD iPSC lines used in this study.

It has been suggested that GABAergic neurotransmission plays a very important role in AD pathogenesis such as Aβ toxicity, hyperphosphorylation of tau and APOE effect (Burgos-Ramos et al., 2008; Jo et al., 2014; Levenga et al., 2013). In light of this information,we modified a previously established embryoid body-based protocol (Liu et al., 2013) to generate iPSC-derived neuronal cultures enriched in GABAergic interneurons. Fig. 1A shows the timeline schematic for the protocol, in which all iPSC lines were successfully differentiated into neuronal networks enriched in GABAergic interneurons in a course of ∼80 days (Supplementary Fig.1A). To qualitatively characterize the progression of differentiation, we performed immunostaining for various markers through the differentiation process. Neural rosettes expressed the progenitor markers PAX6 and Nestin (Fig. 1B) and after being selected and grown as neurospheres for 7 days, the progenitor cells (SOX1^+^) acquired predominantly a forebrain identity due to the expression of the medial ganglionic eminence (MGE) transcription factor NKX2.1 (Fig.1C), in addition to the telencephalic transcription factor FOXG1 (Fig. 1D). After maturation, the neural cultures were composed of GFAP^+^ glia cells and neurons expressing the pan-neuronal markers βIII-Tubulin and MAP2 (Fig.1E). Neurons differentiated for ∼80 days expressed the maturation markers Synapsin I (SYN1) and neurofilaments (SMI-32) (Fig.1F), as well as the neurotransmitter-GABA (Fig.1G). In order to assess the maturation status of the neuronal cultures, we performed RNA sequencing to analyze the transcriptome profile at day 80. Fig.1H shows a heat map of Pearson correlation analysis for key maturation neuronal markers together with the glia markers OLIG2 and GFAP in the iPSC-derived neuronal cultures compared to commercially bought RNA from fetal, adult and AD brain. All iPSC-derived neuronal cultures expressed similar levels of dopaminergic and serotonergic markers and higher levels of GABAergic interneuron markers. To complement and independently confirm these expression data, quantitative real-time PCR (qRT-PCR) analysis was carried out to evaluate the expression levels of GABAergic interneuron markers *PV, SOM, CALB2, GAD67 and GAD65* (Supplementary Fig.1E). Despite the variability of expression levels of the different markers, we observed that the iPSC-derived neuronal cultures might be composed mostly of somatostain (SST) and calretinin (CALB2) subtypes of GABA interneurons. Moreover, due to the low expression level observed for *TREM2* when compared with the commercially bought fetal, adult and AD brain RNA, qRT-PCR was performed for all iPSC-derived neuronal cultures. *TREM2* is expressed in all lines but however significantly upregulated in AD TREM2-2 (Fig.1I). Taken together, we propose (i) the expression of the *TREM2* R47H variant in the AD TREM2-2 and AD TREM2-4 lines has no significant effect on the neuronal differentiation capacity when compared to the control lines CON8 and CON9, as assayed by global transcriptome analysis and RT-PCR. (ii) Though we have not analysed our neuronal cultures for the presence of microglia, probably the mixed neuronal culture might have harbored these.

**Fig 1.**
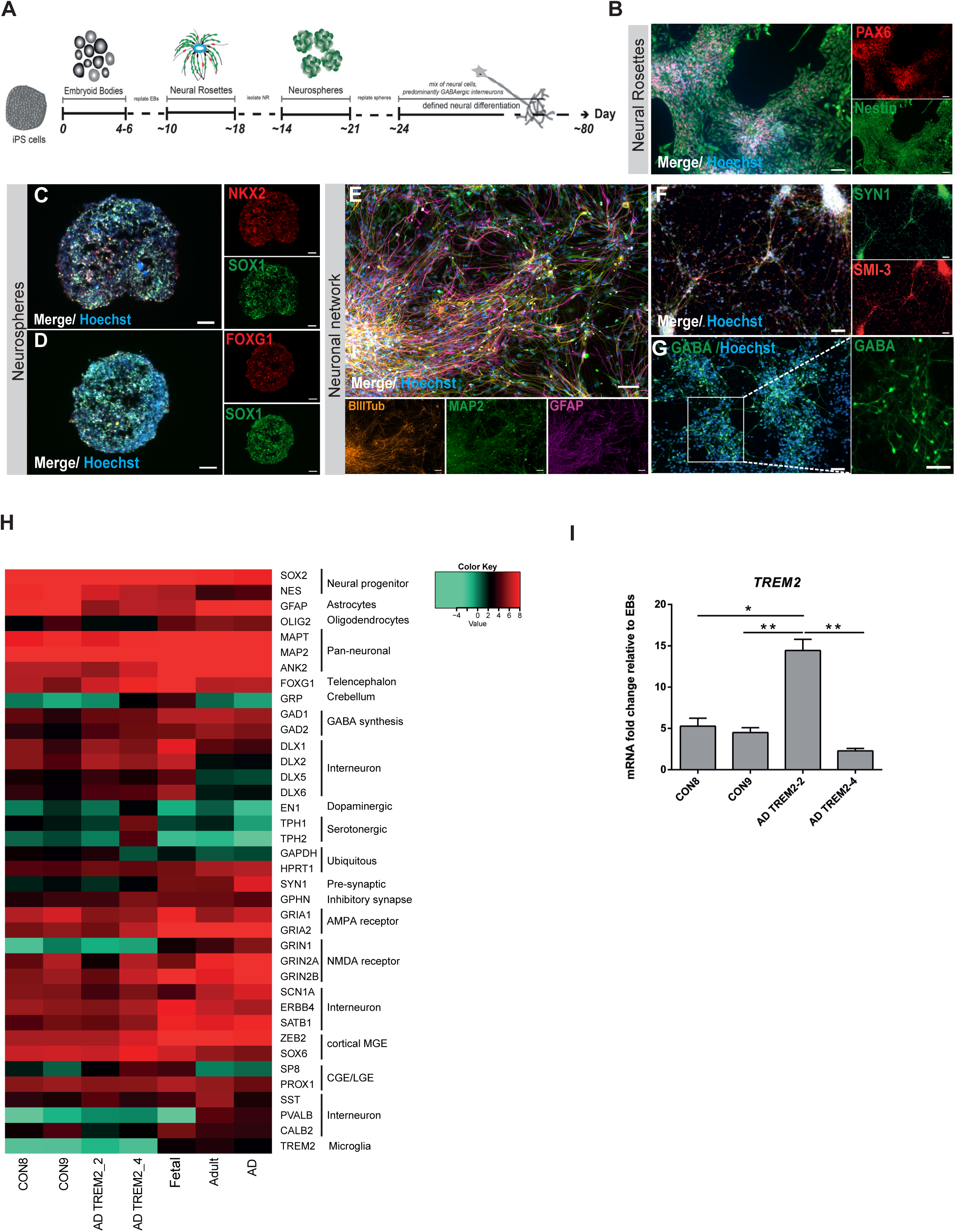
Differentiation and characterization of iPSC-derived neuronal cultures. **A.** Scheme illustrating the main stages of the differentiation protocol for generating iPSC-derived neuronal network enriched in GABAergic interneurons. **B-G.** Representative immunocytochemistry images of **(B)** Neural Rosettes expressing the progenitor markers PAX6 (red) and Nestin (green), **(C)** Neurosphere expressing the progenitor marker SOX1 (green) and the MGE marker NKX2.1 (red), **(D)** Neurosphere expressing the progenitor marker SOX1 (green) and the forebrain marker FOXG1 (red), **(E)** Neuronal network expressing the pan-neuronal markers βIIITub (orange) and MAP2 (green) as well as GFAP (magenta), **(F)** Neural maturation markers SYN1 (green) and SMI-3 (red) and **(G)** Interneurons expressing the neurotransmitter GABA (green). Nuclei are stained with Hoechst. Scale bar, 50 µm. **H.** Heat map of Pearson correlation analysis of RNA-seq data from neural differentiation of control (CON8 and CON9) and AD lines (AD TREM2_2 and AD TREM2_4) and commercially bought RNA from fetal, adult and AD brain for neural progenitor, early neuronal and mature dopaminergic, serotonergic, GABAergic interneuronal markers and glia markers. **I.** Relative gene expression of *TREM2* in iPSC-derived GABAergic interneurons network from control and AD lines shown as fold change to. *p<0.05, ** p<0.01, one-way ANOVA, followed by Tukeýs multiple comparisons test. Data are presented as mean ± SEM from three independent experiments.

### AD TREM2 neuronal network shows a distinct gene expression associated with metabolism and immune-related pathways

To obtain a global view of the transcriptome changes between the AD (AD TREM2-2 and AD TREM2-4) and the control (CON8 and CON9) neuronal cultures, we performed a screening for differential expressed genes (DEGs). Employing RNA-seq we identified 4990 genes exclusively expressed in the AD TREM2 neuronal network (Fig. 2A). BiNGO was used to perform GO term enrichment analysis of the 4990 genes, the results are illustrated as a tree-like structure (Fig. 2B, Supplementary Table 1). In depth analyses of the cellular component identified, significant enrichment associated with *membrane* and *extracellular space*. With regard to biological processes, these proteins were significantly enriched in processes related to *response to stimulus* and *transport*. Moreover, molecular functions such as *signal transducer activity, receptor activity* and *transporter activity,* including *ion membrane transporter activity* and *channel activity* were significantly enriched. KEGG pathway analysis revealed metabolic pathways, which include *drug metabolism – cytochrome P450*, *retinol metabolism,* and *steroid hormone biosynthesis* together with *neuroactive ligand-receptor interactor* (Fig. 1C). As it has been shown that TREM2 regulates innate immunity in AD (Li and Zhang, 2018), we additionally analysed GO terms for biological processes of immune-related genes within the 4990 gene set. Remarkably, 14 significantly enriched terms associated with the regulation of innate and adaptive immune response were identified (Fig. 1D). Overall, these data may suggest that AD TREM2 lines exhibit alterations in key signaling pathways related to metabolism and immune system.

**Fig 2.**
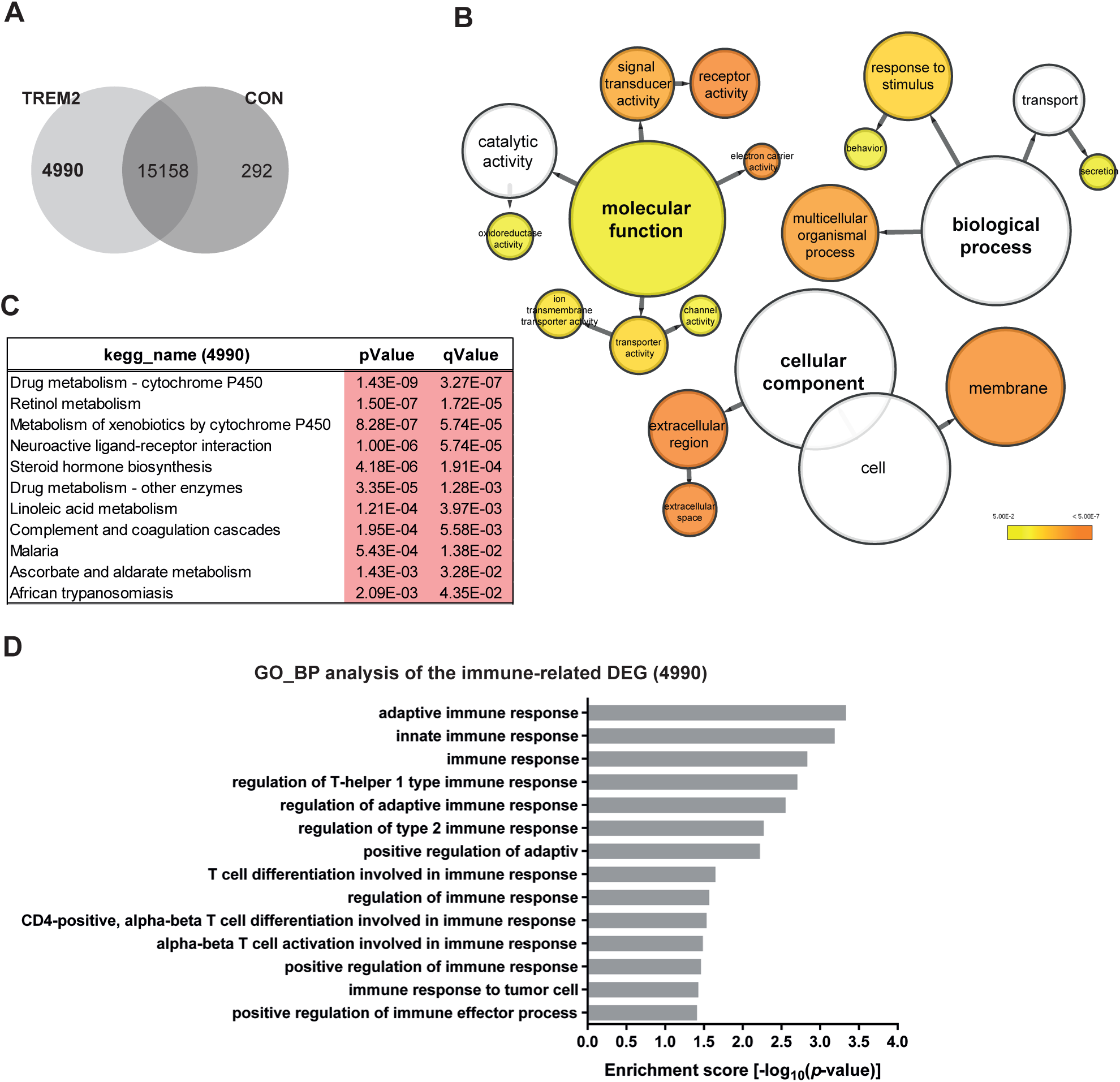
Distinct gene expression profiles in TREM2 neural networks. **A.** Venn diagram illustrating genes exclusively expressed in the TREM2 neural network (4990), the CON network (292) or common between both (intersection-15158) (detection *p* value < 0.05). **B.** BiNGO analysis of the DEGs (4990) exclusively expressed in the TREM2 neuronal network (4990). The orange color of the circles correspond to the level of significance of the over-represented GO category and the size of the circles is proportional to the number of genes in the category (*p*-value <0.05). **C.** KEGG pathway analysis of the genes exclusively expressed in the TREM2 neuronal network (4990). **D.** Significantly enriched gene ontology (GO) terms (Biological Processes) of the genes exclusively expressed in the TREM2 neuronal network (4990) associated with immune system processes (*p*-value <0.05).

### Characterization of AD hallmarks in CON and AD TREM2 neuronal cultures

Numerous evidence support the notion that the small oligomers of Aβ_42_ are intricately associated with the amyloid cascade (Sengupta et al., 2016; Viola and Klein, 2015). However, recent studies have shown that Aβ dimers, abundantly detected in brains of AD patients, are sufficient to account for neurotoxicity and initiating the amyloid cascade (Brinkmalm et al., 2019; Klyubin et al., 2008; Mc Donald et al., 2010; Shankar et al., 2008). Here, we aimed at investigating the effects of the *TREM2* R47H mutation in Aβ production as well as the response of CON and AD iPSC-derived neuronal cultures to stimulations with the well described Aβ-S8C dimer (Abdel-Hafiz et al., 2018; Müller-Schiffmann et al., 2011; Müller-Schiffmann et al., 2016). After 4 months of differentiation, neurospheres were dissociated into single cells and further differentiated 6 weeks. Aβ levels were measured and the neuronal cultures were stimulated with 500 nM of the Aβ-S8C dimer for 72h (Fig. 3A). Conditioned media from the non-stimulated CON and AD lines were analyzed for comparative Aβ_40_ and Aβ_42_ levels employing ELISA. Interestingly, neurons derived from the AD iPSCs lines (AD TREM2-2 and AD TREM2-4) and the CON iPSCs lines (CON8 and CON9) secreted Aβ with similar Aβ_42_ ratio (Fig. 3B-D). We further performed Western Blot analysis to evaluate the levels of tau phosphorylation at Ser202/Thr205 (AT8 epitope), total tau and total APP after stimulation with the Aβ-S8C dimer (Fig. 3E). Although phosphorylation of tau was found in all neuronal cultures, no significant differences in the expression levels of total tau (Fig. 3F) and phosphorylated tau (Fig. 3G) were observed between AD and CON lines after treatment with the Aβ-S8C dimer. Surprisingly, the results revealed that stimulation with the Aβ-S8C dimer induced a modest and uniform increase in the expression levels of APP in all CON and AD neuronal cultures (Fig. 3H). To focus on the effect of the Aβ-S8C dimer, we performed the quantification of APP levels in pooled samples, and this revealed significantly increased APP expression (Fig. 3I). Taken together, these results confirmed that the neuronal cultures (CON8 and AD) secrete Aβ and although no significant differences in the expression of total and phosphorylated tau were observed, APP expression was significantly elevated after Aβ-S8C dimer stimulation. We therefore conclude that the CON and the TREM2 R47H AD iPSC-derived neuronal cultures were capable of recapitulating *in vitro* the hallmarks of AD-like cellular pathology.

**Fig 3.**
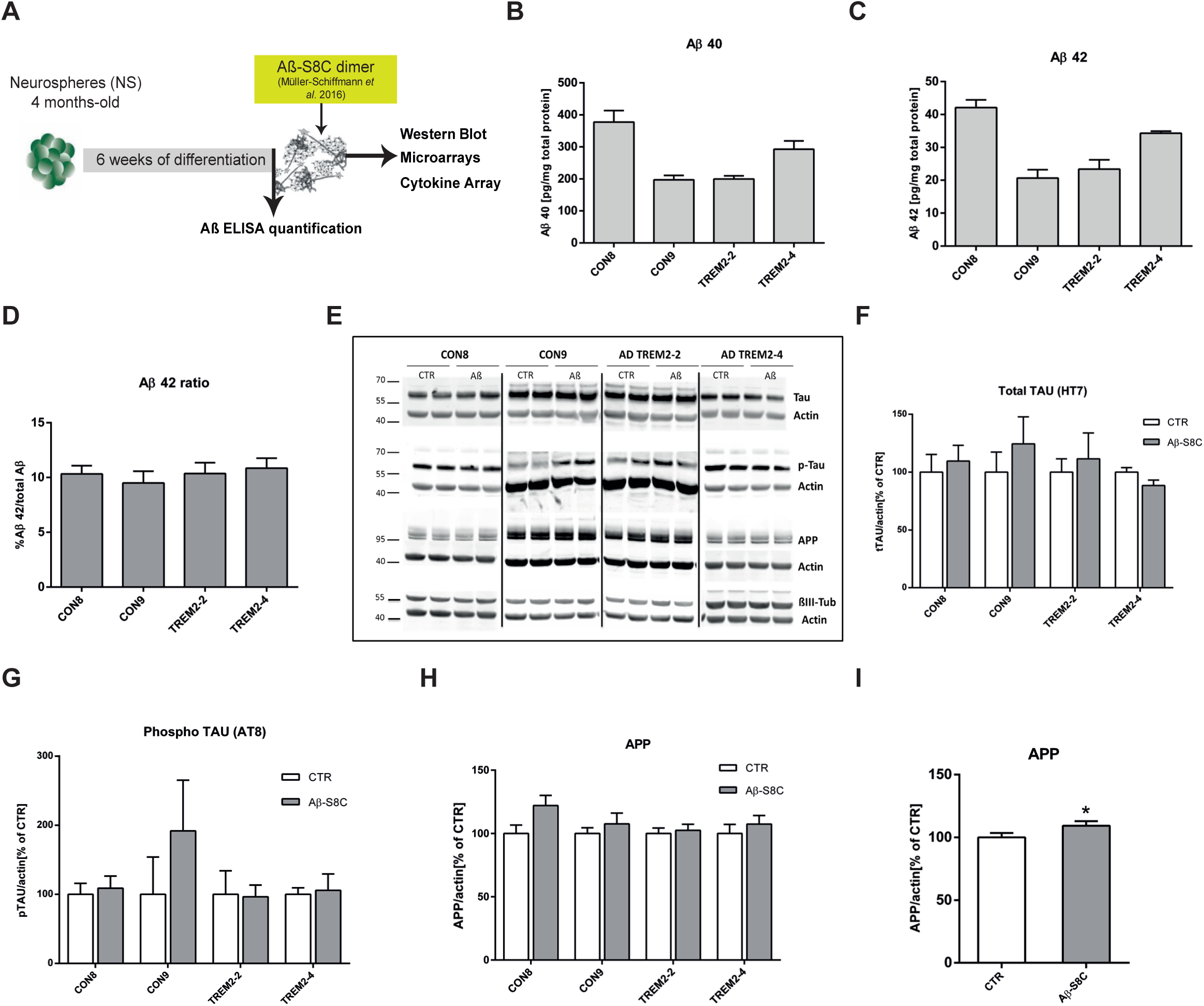
Stimulation of iPSC-derived neuronal cultures with the Aβ-S8C dimer. **A.** Scheme illustrating the approach. Neurospheres were maintained for 4 months in culture, dissociated into single cells, differentiated for 6 weeks then stimulated with 500 nM of Aβ-S8C dimer for 72h. Western blotting, microarrays and cytokine arrays were performed. **B-D.** ELISA quantification of **(B)** total Aβ_40_, **(C)** total Aβ_42_ levels and **(D)** Aβ_42_/Aβ_40+42_ ratio from media collected from the interneuronal network and normalized to the total protein content. All data are presented as mean ± SEM from six independent experiments. **E.** Representative Western blot images of endogenous tau, phosphorylated tau (Ser 202 and Thr 205), APP and the neural differentiation marker βIII-Tubulin after stimulation with 500 nM of the Aβ-S8C dimer. β-ACTIN was used as a loading control. **F-H.** Quantification of **(F)** Total tau, **(G)** phosphorylated tau and **(H)** APP levels. Results are normalized against β-ACTIN and shown as a percentage of control (CTR). All data are presented as mean ± SEM from 3 independent experiments. **I.** Effect of the Aβ-S8C dimer on APP levels in iPSCs derived neuronal network (CON8, CON9, AD TREM2-2 and AD TREM2-4) compared to control. Data are presented as mean ± SEM from 3 independent experiments from 4 biological replicates. *p<0.05, one-tail t-test versus control.

### The Aβ-S8C dimer induces metabolic dysregulation in AD TREM2 neuronal network

To assess the impact of the Aβ-S8C dimer on the gene expression profile of CON and AD iPSC-derived neuronal cultures, we performed transcriptome analysis of CON8 and AD TREM2-4 iPSC-derived neuronal network before and post stimulation with the Aβ-S8C dimer. This analysis identified differential expressed genes (DEGs) between control and Aβ-S8C dimer treatment. Hierarchical cluster analysis revealed a clear separation of CON8 and AD-TREM2-4 iPSC-derived neuronal cultures (Fig. 4A). Remarkably, CON8_ Aβ clustered separately from TREM2-4_ Aβ, therefore implying that genetic background effects were more pronounced than the response elicited by the Aβ-S8C dimer.

**Fig 4.**
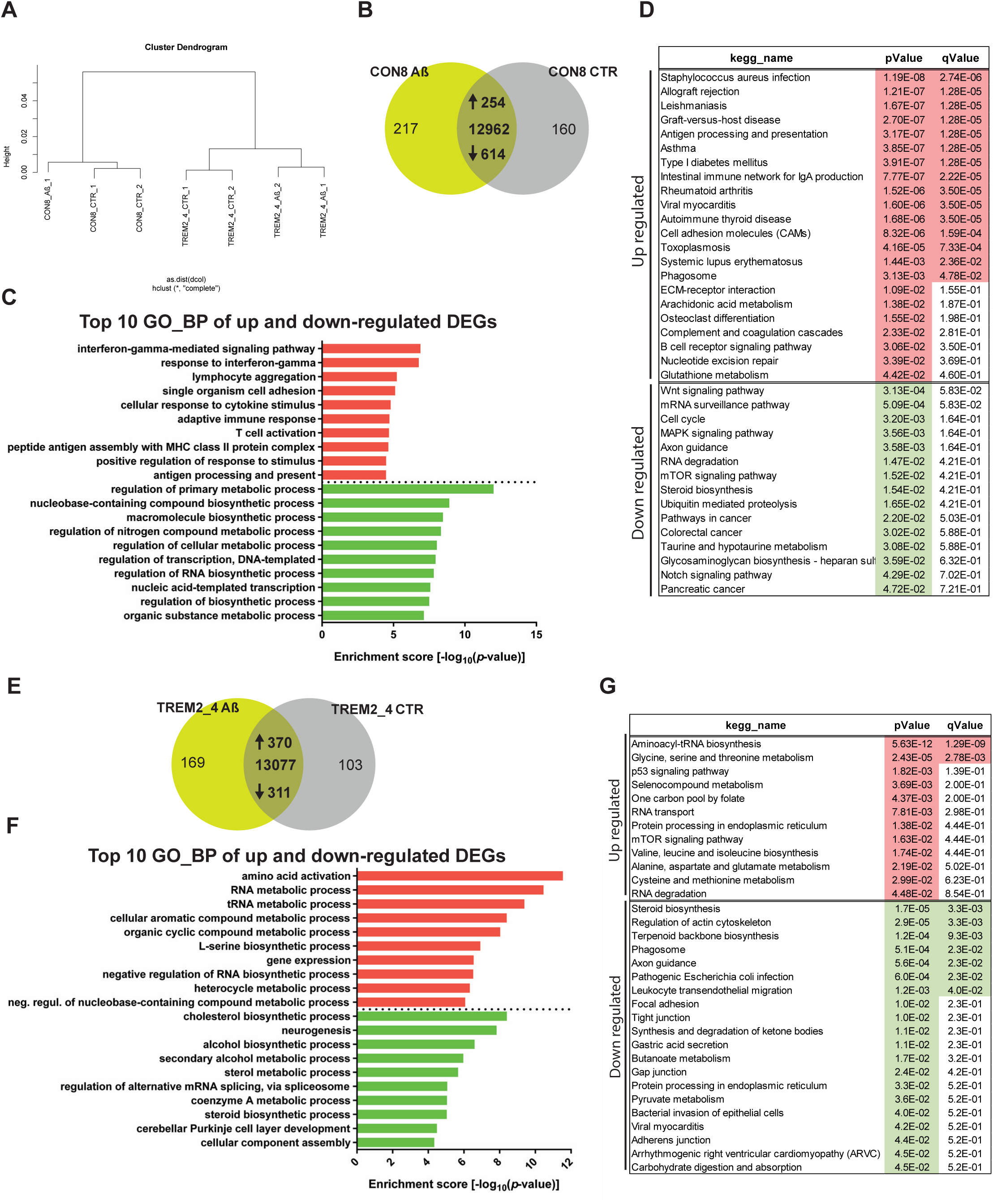
Gene expression profiles of the CON8 and TREM2 neural network stimulated with the Aβ-S8C dimer. **A.** Dendogram obtained by hierarchical cluster analysis of microarray-based (Affymetrix) gene expression data for CON8 and TREM2-4 stimulated with the Aβ-S8C peptide. Transcriptomes of CON8_CTR cluster with CON8_ Aβ while those of the TREM2-4_CTR cluster separately with TREM2-4_ Aβ. **B**. Venn diagram showing genes expressed only in the CON8 neural network subjected to Aβ-S8C peptide stimulation (green), the control condition (grey), and common to both conditions (intersection) (detection *p* value < 0.05). **C.** Top 10 significantly enriched gene ontology (GO) terms for Biological Processes of DEGs upregulated (254-red) and down –regulated (614-green) subjected to Aβ-S8C peptide stimulation in CON8 neural network (*p*-value <0.05). D. KEGG enrichment analysis of up- and downregulated DEGs (*p*-value <0.05). **D.** Venn diagram showing genes expressed only in the AD TREM2_4 neural network when stimulated with the Aβ-S8C peptide (green), the control condition (grey), and common to both conditions (intersection) (detection *p* value < 0.05). **F.** Top 10 significantly enriched gene ontology (GO) terms for Biological Processes of DEGs upregulated (370-red) and down –regulated (311-green) subjected to Aβ-S8C peptide stimulation in the AD TREM2_4 neural network (*p*-value <0.05). **G.** KEGG enrichment analysis of up- and downregulated DEGs (*p*-value <0.05).

Evaluation of DEGs in CON8 neuronal network before and after stimulation with the Aβ-S8C dimer identified 868 genes (Fig. 4B), 254 were upregulated and 614 downregulated (Supplementary Table 1). Figure 4C shows the related Top10 GO BP (biological processes) terms. Upregulated genes in CON8_Aβ were significantly enriched for GO terms such as *interferon-gamma-mediated signaling pathway* and *cellular response to cytokine stimulus*. In contrast, the downregulated genes in CON8_ Aβ in comparison to control were associated with the GO terms-*regulation of primary metabolic processes* and *regulation of RNA biosynthetic process*. In agreement with the GO analysis, KEGG pathway analysis for the same set of genes revealed the upregulated genes in CON8_ Aβ to be involved in pathways related to inflammatory responses, for example, *staphylococcus aureus infection* and *antigen processing and presentation*. In addition, CON8_ Aβ also showed upregulation of the *phagosome* pathway, while *Wnt signaling pathway* and *axon guidance* were among the downregulated KEGG pathways (Fig. 4D, Supplementary Table 1).

Focusing on *TREM2* R47H, 681 DEGs were identified when comparing Aβ-S8C dimer stimulated and non-stimulated AD TREM2-4 neuronal cultures. (Fig. 4F), of these 370 were upregulated and 311 downregulated (Supplementary Table 2). Figure 4F lists the Top10 GO BP terms. The upregulated genes in AD TREM2-4_ Aβ were significantly enriched for *amino acid activation* and *RNA metabolic process*. In contrast, the downregulated genes were associated amongst others with *cholesterol biosynthetic process* and *neurogenesis*. KEGG pathway analysis revealed upregulation of pathways such as *glycine, serine and threonine metabolism, p53 signaling pathway and mTOR signaling pathway.* Surprisingly, in contrast to CON8_Aβ, AD TREM2-4 neuronal cultures stimulated with Aβ-S8C dimers showed down-regulation of the *phagosome* pathway (Fig. 4G, Supplementary Table 2). Taken together, these results revealed that the AD TREM2-4 neuronal cultures respond in a unique way to Aβ-S8C dimer stimulation, namely a metabolic dysregulation in contrast to an inflammatory response, which could be observed in the CON8 neuronal cultures.

### Aβ-S8C dimer stimulation of the AD TREM2 neuronal network revealed indications of impaired phagocytosis-related pathway

TREM2 is crucial for regulating phagocytosis in microglia and although the effect in phagocytosis by the AD-associated *TREM2* mutations have recently been a focus of study, it remains unclear if *TREM2* R47H variant indeed impairs phagocytosis (Brownjohn et al., 2018; Claes et al., 2018; Garcia-Reitboeck et al., 2018; Hsieh et al., 2009; Kleinberger et al., 2014; Takahashi et al., 2005). As described above, *phagocytosis* appeared as significantly upregulated pathway in CON8 but was downregulated in AD TREM2-4 neuronal cultures after Aβ stimulation. We then analyzed differential expression of genes involved in this pathway. Figure 5 depicts the KEGG annotated *phagosome pathway* with upregulated genes in CON8_ Aβ colored in red and those downregulated in AD TREM2-4_ Aβ colored in green. After stimulation with the Aβ-S8C dimer, CON8 revealed up-regulation of *HLA-DMA*, *HLA-DMB. HLA-DOA, HLA-DPB1, HLA-DQA1, HLA-DQB1, HLA-DRB1, and HLA-F*, all genes belonging to the Major Histocompatibility complex II (MHCII). In contrast to the AD TREM2-4 non-stimulated networks, stimulation with the Aβ-S8C dimer induced down-regulation of *TUBB4A, TUBB4B, DYNC1H1, LAMP2, ATP6V1A, ACTB, THBS1, CALR and TUBBA1C*. Supplementary Table 3 shows the relative mRNA expression. Taken together, these results suggest that neuronal cultures harboring the *TREM2* R47H variant but not controls likely undergo an impaired phagocytosis response in the presence of the Aβ-S8C dimer.

**Fig 5.**
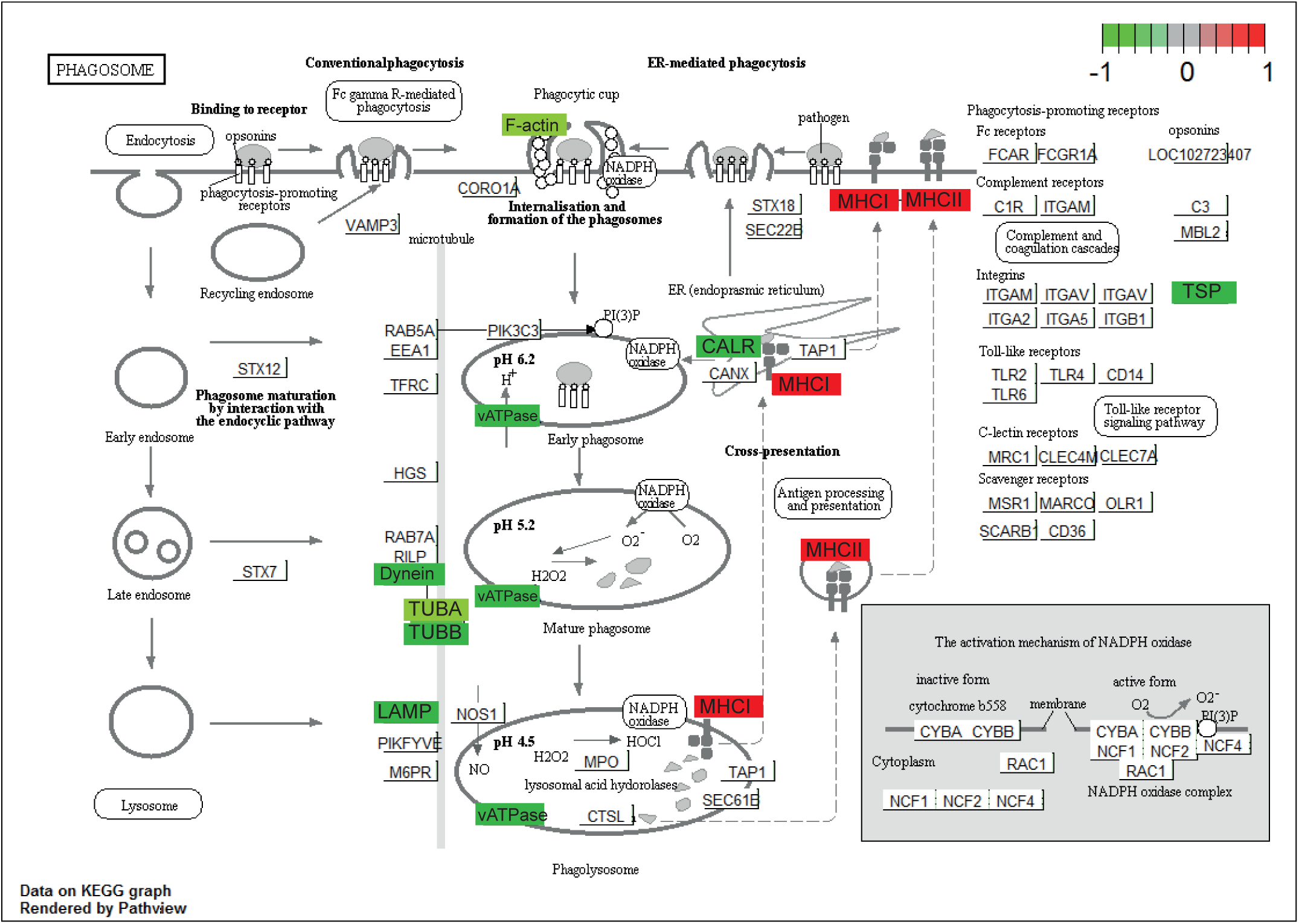
Representation of the KEGG Phagosome Pathway. Upregulated DEGs genes in response to Aβ-S8C peptide stimulation in CON8 neuronal cultures are shown as red boxes and downregulated DEGs in TREM2-4 neuronal cultures are shown as green boxes.

### TREM2 R47H neuronal cultures show compromised inflammatory response related gene expression pattern upon stimulation with the Aβ-S8C dimer

Based on the fact that dysregulated cytokine production from microglia, astrocytes and neurons are associated with the development of AD (Domingues et al., 2017), we analyzed the cytokine expression profile as well as the secretion profile from the AD TREM2 neuronal cultures after stimulation with the Aβ-S8C dimer. Employing microarray-based global gene expression data, a heatmap-based analysis of 100 key cytokines (extracted from the Proteome Profiler Human XL Cytokine Array, R&D systems) revealed that stimulation with the Aβ-S8C dimer induced transcriptional changes for several of these genes in AD-TREM2-4 (Fig.6A). Interestingly, the AD TREM2-2 neuronal network showed down-regulation of cytokines, chemokines and acute phase genes such as *IL1RL1, IL13, IL15, IL16, IL27, IL32, CXCL10, CXCL11, TFRC, SERPINE1, C5, THBS1, RLN2, SPP1, EGF, LIF, GC, BSG, MPO, CST3, FLT3LG and CCL20*. Surprisingly, only *IGFBP2, RBP4, VEGFA, CXCL5, IL19 and TDGF1* had higher expression levels after Aβ-S8C dimer stimulation when compared to the control samples. We next aimed at determining if stimulation the Aβ-S8C dimer could also alter the secretion of cytokines and chemokines in the AD TREM2 neuronal cultures. To this end, we collected the cell culture supernatants from the AD TREM2-2 and AD-TREM2-4 neuronal cultures 72h post stimulation with the Aβ-S8C dimer and from non-stimulated controls. Thereafter, we carried out secretome analysis employing the Proteome Profiler Cytokine Array (Fig.6B). In agreement with the previous results, the level of secretion of all cytokines and chemokines decreased after Aβ-S8C dimer stimulation when compared to control (Fig. 6C), with the exception of ICAM-1, MIF and SerpinE1. Taken together, these results would imply that AD TREM2 neuronal cultures compromise the efficient activation of the inflammasome pathway in response to Aβ-S8C dimer stimulation.

**Fig 6.**
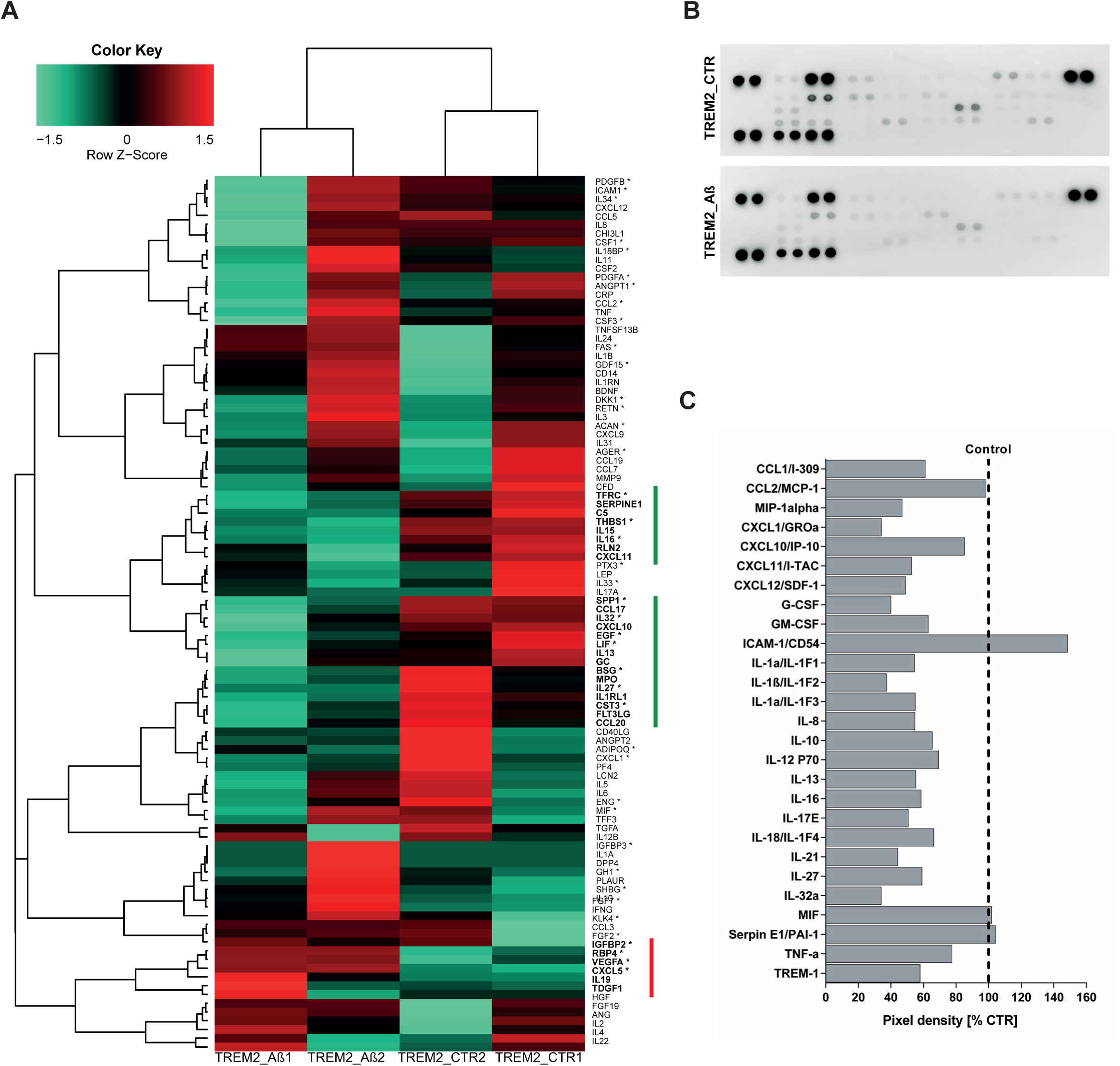
Analysis of cytokine expression and secreted factors in TREM2 neuronal network under Aβ-S8C stimulation. **A.** Heat map of Pearson correlation analysis of microarrays data from TREM2 neural differentiation under Aβ-S8C stimulation (TREM2_ Aβ1 and TREM2_ Aβ2) or control (TREM2_ CTR1 and TREM2_ CTR2) showing the differential expression of cytokines. The highlighted green genes represent a cluster of cytokines downregulated under Aβ-S8C stimulation whereas the highlighted red genes represent a cluster of cytokines upregulated. **B.** Human cytokine array showing the effect of the Aβ-S8C peptide on the secreted factors of neuronal cultures from pooled AD TREM2-2 and AD TREM2-4 culture supernatants of control condition and 72h of Aβ-S8C stimulation. **C.** Quantitative analysis of the secreted factors shows that Aβ-S8C treatment decreases the amount of secreted cytokines in AD TREM2 neuronal cultures.

### Protein-protein interaction (PPI) network revealed the TREM2 R47H-depended Aβ-S8C signature

To gain insights into the gene expression signature triggered by the Aβ-S8C dimer in LOAD, we focused on the genes exclusively expressed in the AD TREM2 neuronal culture after stimulation with the Aβ-S8C dimer. A Venn diagram analysis showed that most (12687) genes were expressed in common between CON and AD TREM2 lines with and without Aβ-S8C dimer stimulation (Fig. 7A, Supplementary table 4). However, 95 genes were exclusively expressed in AD TREM2 neuronal cultures stimulated with Aβ-S8C dimer. GO analysis (Fig. 7B) delivered several terms related to neuron and immune-system related processes including *stimulatory C-type lectin receptor signaling pathway* as most significant. Pathway analysis of the 95 TREM2-Aβ-S8C genes (Fig. 7C) revealed *Neuroactive ligand-receptor interaction* as most significant pathway and *Metabolic pathways* with the higher number of genes. The 95 genes were further analysed in a protein interaction network (PPI) based on interactions from the BioGrid database resulting in a network containing APP and a big hub centered around *HSPA5* which encoded the endoplasmic reticulum chaperone BiP (Fig. 7D). HSPA5 has been reported to control the activation of the unfolding protein response (UPR), a pro-survival pathway in response to ER stress caused by misfolded proteins. Since there is evidence that the ER stress response, namely the UPR plays a role in the pathogenesis of AD (Doyle et al., 2011), we took a deeper look into the GO terms related to ER after Aβ-S8C dimer stimulation (Supplementary table 2, highlighted in yellow). We observed that the Aβ-S8C dimer triggered a ER stress response, increasing the expression of *ATF3* and *DDIT* in both CON and TREM2. Interestingly, the ER stress response was more prominent in the AD TREM2 neuronal cultures, where several genes from the UPR were up-regulated (*XBP1, AT4, PUMA, HERPUD1*) and *HSPA5* and *CALR* were down-regulated (Fig.8, Supplementary table 5). These results highlight the unique response triggered by the Aβ-S8C dimer in the AD TREM2 neuronal cultures. By creating of a PPI network we were able to link the Aβ-S8C signature genes with the ER-stress, namely the activation of UPR.

**Fig 7.**
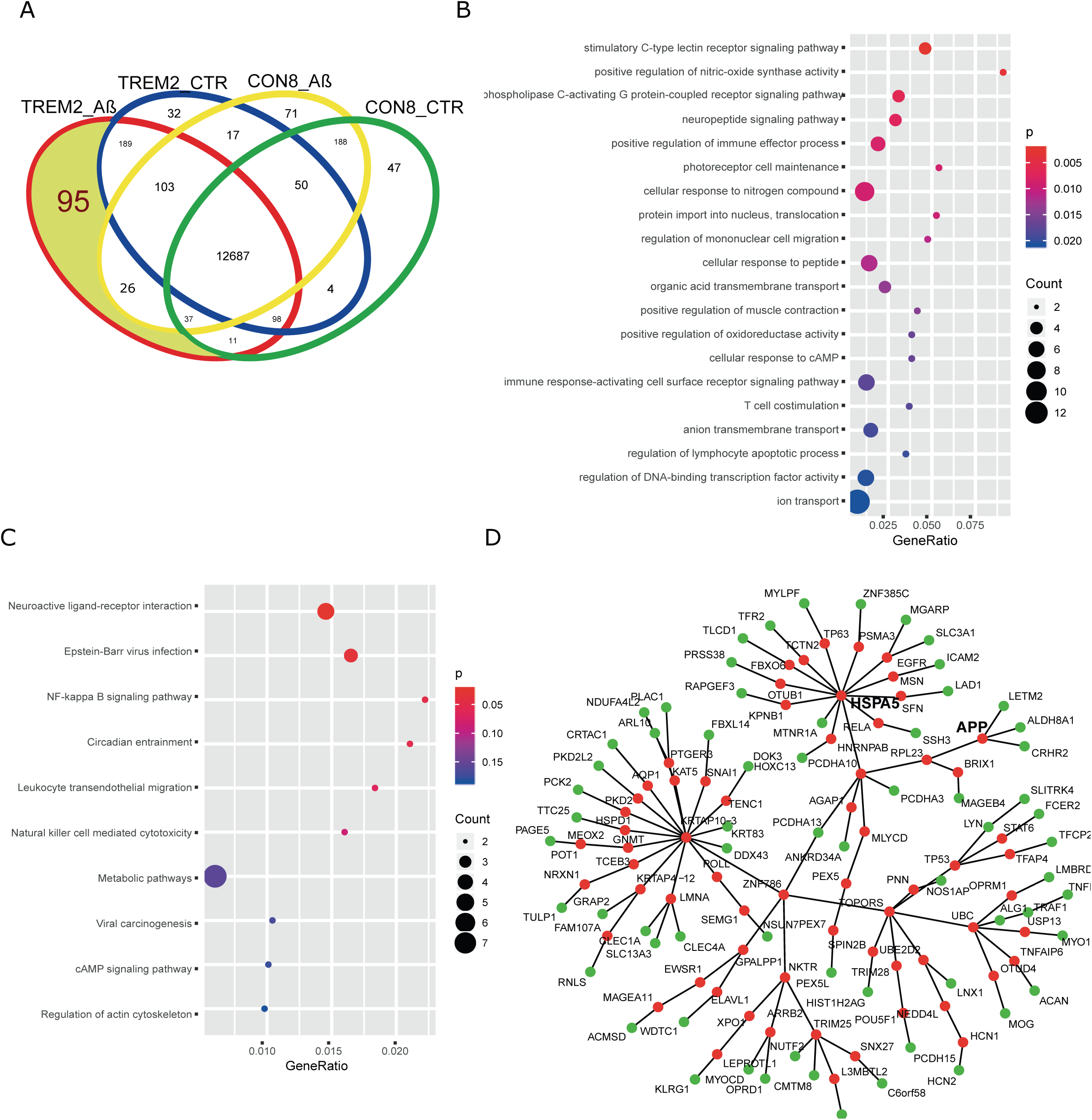
Aβ-S8C stimulated TREM2 neuronal network activate a protein interaction network including APP and HSPA5. **A**. Venn diagram dissecting 95 genes expressed in Aβ-S8C stimulated TREM2 neuronal cultures from genes expressed in TREM2 control and healthy neuronal cultures with and without Aβ-S8C stimulation. **B.** Dot plot of Gene ontologies (Biological Process) overrepresented in the 95 TREM2_Aβ genes **C.** Dot plot of KEGG pathways overrepresented in the 95 TREM2_Aβ genes. **D**. Protein-protein interaction network derived from the 95 TREM2_Aβ genes with APP and HSPA5. Nodes from the 95 genes are colored green and the nodes added using the Biogrid database to connect the network are colored red.

**Fig 8.**
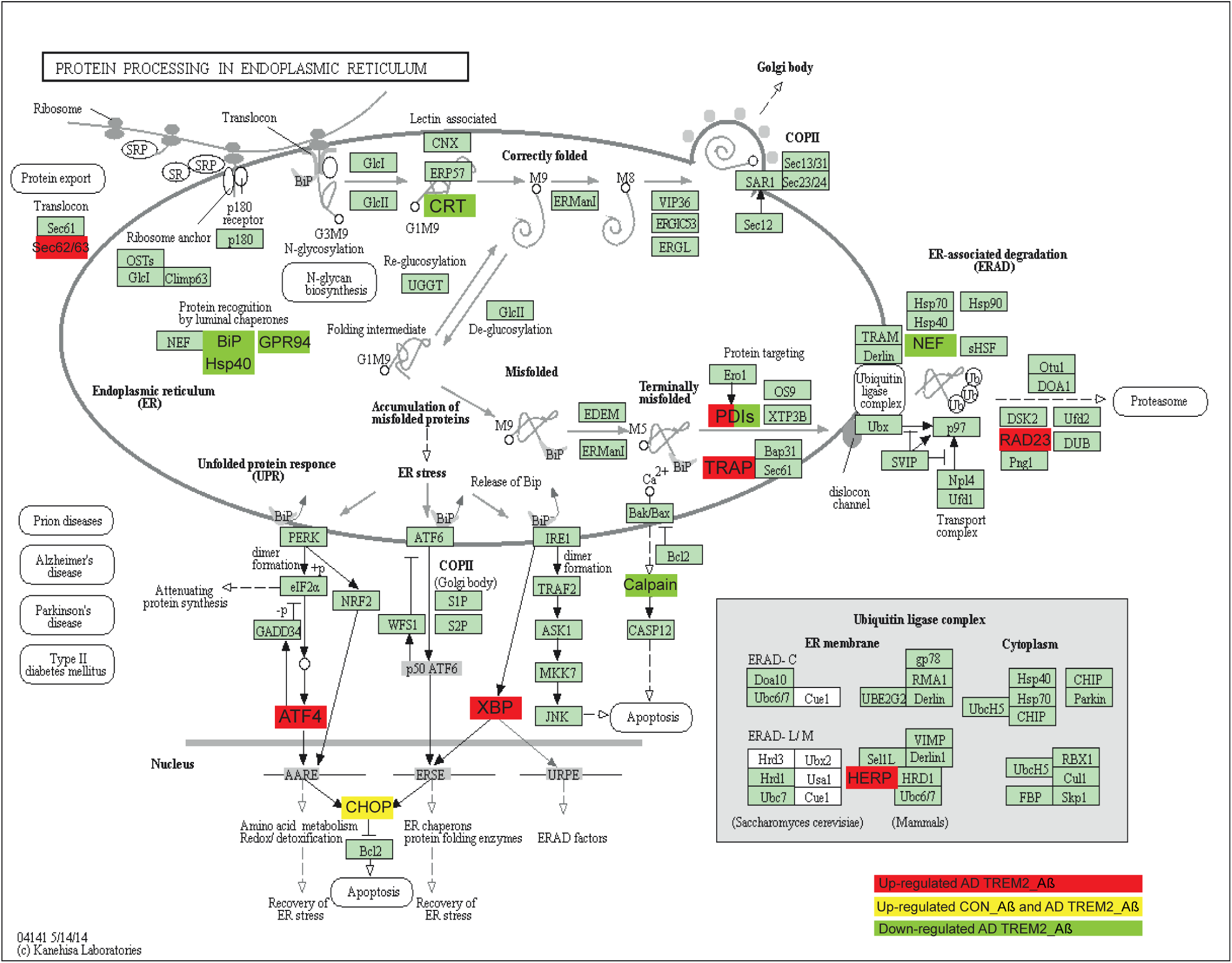
Representation of the KEGG Protein Process in Endoplasmic Reticulum pathway. Upregulated DEGs genes in response to Aβ-S8C stimulation in TREM2-4 neuronal cultures are shown as red boxes, upregulated DEGs genes in response to Aβ-S8C stimulation in CON 8 and TREM2-4 neuronal cultures are shown in yellow boxes and downregulated DEGs in TREM2-4 neuronal cultures are shown as green boxes.

## Discussion

While the mechanisms underlying the etiology of AD have been a focus of study over several decades, the current knowledge about the etiology and pathogenesis of AD are still incomplete. The use of primary neurons from animal models and immortalized cell lines based on modifications in *APP*, *PSEN1* and *PSEN2* has provided some insights into EOAD. While these models are helpful for studying a specific causal mutation (EOAD), there are several hurdles and limitations associated with studying LOAD, which requires the endogenous expression of genetic mutations and their genetic interactions. Understanding the biological implications of the recently identified genetic risk variants, namely the R47H substitution in*TREM2,* is essential to enable the establishment of genotype-phenotype correlations, which can lead to potential novel therapeutic approaches. The breakthrough development of iPSCs technology provides the most applicable tool to create an *in vitro* sporadic patient-derived model. Although modeling AD using patient-derived iPSCs has been prominent, a handful of studies to date have generated and characterized iPSC-derived neuronal cultures from LOAD patients (Flamier et al., 2018; Hossini et al., 2015; Israel et al., 2012; Muratore et al., 2014; Ochalek et al., 2017). This is the first study describing the generation and characterization of a model of LOAD based on Aβ dimer stimulated neuronal network originating from lymphoblast-derived iPSCs derived from LOAD patients carrying the missense mutation R47H in *TREM2*.

First, we differentiated the iPSCs to neurons using a modified protocol described by Liu *et al*., 2013 and analyzed the distinct progression steps during the differentiation process. Transcriptome analysis and ICC confirmed the ability of our modified protocol to derive neurons and glia cells within our neuronal culture. Based on gene expression comparison between the iPSC-neuronal cultures with the commercially bought fetal, adult, and AD brain RNA we could show that our cells express the expected maturation markers. Thus, our results imply that lymphoblast-derived iPSCs from LOAD patients and healthy donors can be robustly differentiated into neuronal cultures. Moreover, we did not observe profound difference in the differentiation and maturation propensity between iPSCs derived from LOAD patients and healthy donors-in agreement with previous reports (Flamier et al., 2018; Hossini et al., 2015; Israel et al., 2012; Ochalek et al., 2017). Cheng-Hathaway *et al*., and Sudom *et al.,* reported that *Trem2* R47H knock-in mice showed reduced *Trem2* mRNA and protein expression in the brain as well as reduced soluble fragments of Trem2 (sTrem2) in plasma (Cheng-Hathaway et al., 2018; Sudom et al., 2018). More recently, Xiang *et al*. reported that a mouse-specific splicing caused this reduction and *TREM2* mRNA levels were normal in both iPSC-derived microglia and in patient brains with the *TREM2* R47H variant (Xiang et al., 2018). We therefore evaluated if T*REM2* expression was different in LOAD patients carrying the *TREM2* R47H variant compared to the control. Our results indicate that *TREM2* mRNA was significantly upregulated in the AD TREM2-2 but not in the AD TREM2-4 neuronal cultures compared to the control. This ambiguity is probably due to the limitations imposed by our small sample size. In addition, our neuronal network is composed mainly of neurons so the TREM2 positive cells are in low abundance. In a recent cross-sectional study, Suárez-Calvet et al., 2019 reported that sTREM2 levels in the cerebrospinal fluid (CSF) are higher in patients carrying the R47H variant compared to non-carriers AD patients. Further studies are needed to evaluate the levels of sTREM2 and its relationship with the expression of TREM2 on the surface of the microglia.

Although the neuronal cultures derived from LOAD patients and healthy donors did not exhibit differences in morphology or expression of differentiation markers, transcriptome analysis showed a distinct profile. Interestingly, GO analysis revealed that the proteins encoded by the 4990 genes exclusively expressed in AD TREM2 neuronal cultures were predominantly mapped in the cell membrane and in the extracellular space. These genes were involved in (i) biological processes (BP) terms such as *response to stimulus* and *secretion* and (ii) molecular functions (MF) terms such as *signal transducer activity*, *receptor activity*, *transporter activity*, *channel activity and ion transmembrane transporter activity*. As part of these exclusively expressed genes we also identified genes of the Matrix Metalloproteinases (MMPs) family, for example *MMP2* and *MMP9*. Metalloproteinases play an important role in the pathogenesis of AD. While MMP2 might have a protective role, MMP9 expression, which is increased in AD patients, is induced by Aβ and it can influence tau aggregation (Stomrud et al., 2010). Furthermore, members of the ATP-binding cassette (ABC) and the solute carrier (SLC) families were over-represented in the GO_MF. ABC transporters have been implicated in AD pathophysiology, associated with processes leading to the accumulation of Aβ in the CNS. Importantly, we observed the exclusively expression of GLUT4 (*SLC2A4)*, a crucial insulin sensitive glucose transporter upregulated in AD patients, which is responsible for regulating glucose metabolism in neurons (Katsel et al., 2018; Shah et al., 2012). As anticipated, we also identified GO terms related to the regulation of the innate and adaptive immune response as significantly enriched. Implications of these results are that our AD TREM2 neuronal cultures show a distinct signal transducer and transporter activity that may contribute to metabolic alterations, to an inadequate immune response and ultimately to neurotoxicity. According to the amyloid cascade theory, accumulation of Aβ plays a key role in triggering the cascade of events underlying the pathogenesis of EOAD. However some studies have shown that Aβ secretion is not altered in LOAD-derived neurons (Kondo et al., 2013; Ochalek et al., 2017). In accordance, our results show that AD-TREM2 cultures secreted Aβ with a similar Aβ_1-42_ to Aβ_1-40_ ratio as the control. Nonetheless, the levels of Aβ_1-40_ and Aβ_1-42_ were highly reproducible across multiple differentiations (six) and lines (four), thus establishing our cell culture model as robust for manipulating the production of Aβ. We subsequently aimed at evaluating the potential effect of Aβ in our neuronal cultures in order to close the gap in our understanding of the mechanisms that are underlying the early stages of AD. Aβ-S8C dimer can induce neurotoxicity and abnormal synaptic signaling, together with impaired cognitive functions in the absence of plaque pathology, thus mimicking the early stages of AD (Müller-Schiffmann et al., 2016). Aβ-induced tau hyper-phosphorylation has been described to initiate the signalling cascade alterations that culminate in NFT formation and neuronal degeneration (Wada et al., 1998). Phosphorylation of the AT8 epitope (Ser202/Thr205) has been found to be elevated in sAD-derived neurons (Ochalek et al., 2017). We were not able to detect an increase in phosphorylation of tau at Ser202/Thr205 after Aβ-S8C dimer treatment. In addition, there were no differences in the levels of phosphorylation detected between control and AD TREM2 neuronal cultures in the non-stimulated conditions. Tau can be phosphorylated on more than 80 residues, and it is known that Ser422 is phosphorylated earlier than Ser202/Thr205 during NFTs formation (Cruchaga et al., 2013). Based on these facts and the results obtained, we can assume that duration of incubation of the Aβ-S8C dimer was presumably not enough to detect increase phosphorylation at Ser202/Thr205. Our results show that independent of the genetic background, incubation with the Aβ-S8C dimer increased the levels of total APP. A more in-depth analysis of APP processing will provide more insights into the pathogenic role of the *TREM2* R47H variant in EOAD.

In addition to interfering with total APP levels, Aβ-S8C dimer stimulation induced a remarkable and significant transcriptome change in the control as well as in the AD TREM2 neuronal cultures. Annotation and enrichment analysis revealed that the upregulated DEGs induced by Aβ-S8C stimulation in the control neuronal cultures are related to immune system activation (*interferon-gamma-mediated signaling pathway*, *cellular response to cytokine stimulus* and *adaptive immune response*). Aβ soluble species have also been linked to an attenuation of the Wnt signaling pathway, in addition to putative effects on cell cycle, contributing to synaptic dysfunction and neurodegeneration. In accord, our data showed that the Wnt signaling pathway and cell cycle were downregulated after Aβ-S8C stimulation in the control neuronal cultures. On the contrary, the TREM2 R47H AD neuronal cultures responded in a completely different manner to stimulation with the Aβ-S8C dimer.

The effect of the AD-associated TREM2 mutations on phagocytosis is an active area of study but so far variable results have been obtained. While R47H transduced HEK cells displayed a reduced up-take of latex beads and Aβ_1-42_, no changes were observed in the fluorescent pH-sensitive rhodamine *Escherichia coli* (pHrodo-linked *E.coli)* uptake assay (Kleinberger et al., 2014). Additionally, *TREM2^+/R47H^* transdifferentiated microglia-like cells (Claes et al., 2018) and microglia-like cells derived from *TREM2* T66M ^+/-^, T66M ^-/-^, and W50C^-/-^ hPSCs, also showed no defects in the *E.coli* uptake (Brownjohn et al., 2018; Garcia-Reitboeck et al., 2018). We found that the control neuronal cultures upregulated the phagosome pathway after Aβ-S8C stimulation, namely the genes associated with MHCII. These observations are in line with previous reports where incubation with Aβ led to an accumulation of MHC-II and AD patients also showed up-regulation of MHC-II (Schetters et al., 2017). On the contrary, these genes were not differentially regulated in our AD TREM2 neuronal cultures, but interestingly other genes associated with the phagosome pathway were downregulated. Calreticulin is encoded by the *CALR* gene. It is an endoplasmic reticulum protein that interacts with Aβ, and is considered as a scavenger for Aβ_1-42_ (Duus et al., 2008). Low levels of calreticulin have been observed in AD brains, and it has been suggested that this down-regulation can lead to the pathological processes of AD (Lin et al., 2014). Notably, the levels of tubulins *TUBB4A* and *TUBB4B* were downregulated, supported by Hondius et al., 2016 where the levels of these tubulins identified by mass spectrometry analysis in human post-mortem brain tissue were significantly decreased over the progressive stages of AD. On the other hand, lysosome-associated membrane protein 2 (LAMP-2) together with other lysosome-related proteins was found to be increased in CSF from AD patients (Armstrong et al., 2014). Interestingly, *LAMP2* was downregulated in our AD TREM2 neuronal network leading us to speculate that R47H AD carriers have a unique response to phagocytosis, probably due to the partial loss of function of TREM2 activity.

The analysis of pro-inflammatory cytokines at the levels of mRNA and the secretome of the AD TREM2 neuronal cultures in response to Aβ-S8C stimulation are of particular interest. Although the mRNA expression of the *IL-1β*, *IL-6*, *TNF-α* and *MIP-1α* proinflammatory cytokines was upregulated in some of the samples, the secretion of these cytokines was downregulated. A recent study using iPSC-derived microglia-like cells from patients carrying the T66M and W50C missense mutation within *TREM2* showed that these cells have a deficit in cytokine release (Garcia-Reitboeck et al., 2018). Indeed, *SPP1* and *GPNMB*, encoding osteopontin and osteoactivin, were also downregulated in AD-TREM2 neuronal cultures and not in the control after Aβ-S8C stimulation. SPP1 and GPNMB are microglia activation-related transcripts that are upregulated in AD models and associated with Aβ accumulation. In support of our data, it was recently reported that SPP1 and GPNMB reflect TREM2 signaling and the expression is highly sensitive to the R47H variant (Song et al., 2018). Interestingly, there was a cluster of genes associated with insulin resistance, which was upregulated after Aβ-S8C stimulation. Increased levels of RBP4 were found in APP/PSEN1 mice and in insulin resistant humans (Mody et al., 2011). Along the same track, it has been suggested that IGFBP2 plays a role in AD progression (Bonham et al., 2018). Both of these genes were indeed upregulated in response to Aβ-S8C in our AD TREM2 neuronal cultures, thus further lending credence to the fact that metabolic dysfunction plays an important role in the pathogenesis of AD. It is noticeable that Aβ-S8C triggers a unique response in AD TREM2 neuronal cultures, when compared to control. The creation of a PPI network between the exclusively expressed genes in the AD TREM2 after Aβ-S8C stimulation revealed *HSPA5* as the core of the Aβ-S8C signature. HSPA5, a chaperone protein that upon accumulation of unfolded proteins controls the activation of the UPR sensors (Doyle et al., 2011), was found down-regulated after Aβ-S8C in our AD TREM2 neuronal cultures. Katayama *et al.,* found that HSPA5 levels are reduced in AD patient’s brains. Although Aβ-S8C stimulation up-regulates *ATF3* and *DDIT3* (CHOP) in both CON and AD TREM2 neuronal cultures, the prominent alteration in the UPR was observed in the AD TREM2 cultures with the up-regulation of *XBP1, ATF4, BBC3, HERPUD1* and *CALR.* Supporting our data, several studies have found up-regulation of the UPR in brain samples of AD patients (Hamos et al., 1991; Hoozemans et al., 2005). According to Han *et al,.* insufficient protein-folding homeostasis by URP increases expression of ATF4 and CHOP and initiates the ER-stress-mediated cell death, activating target genes involved in protein synthesis like aminoacyl-tRNA synthetases and RNA metabolic processes leading to oxidative stress and cell death. Interestingly, biological processes related with increased protein synthesis such as amino acid activation and RNA metabolic process together with the KEGG pathway protein processing in the endoplasmic reticulum were up-regulated in AD-TREM2 neuronal cultures. It seems that Aβ-S8C stimulation leads to the activation of the UPR that initially might be protective, however if the balance in the proteostasis is not reestablished, the ER-stress-mediated cell death might mediate the neurodegeneration in AD.

## Conclusion

Our established neuronal cultures using lymphoblast-derived iPSCs from patients harbouring the R47H mutation in *TREM2* is a relevant model for investigating the effect of this variant in the etiology of LOAD. Comparative global transcriptome analysis identified a distinct gene expression profile in AD TREM2 neuronal cultures, further suggesting that these lines exhibit alteration in key signaling pathways related to metabolism and immune system in comparison to control, this implying the partial loss of function of TREM2 due to the R47H substitution. In addition, stimulation with the Aβ-S8C dimer revealed metabolic dysregulation, impaired phagocytosis-related pathway and altered inflammatory responses. Furthermore, our data strongly suggests that the Aβ-S8C dimer signature seems to be centered in the ER-stress response. In conclusion, our AD *TREM2* R47H *in vitro* model is capable of efficiently responding to signaling cascades associated with the AD pathogenesis and thus is a promising cellular tool for investigating the molecular mechanisms underlying LOAD. Additionally, this cellular model can facilitate the discovery of new AD biomarkers, enable toxicology studies as well as the identification of potential drug targets for future therapy of this devastating disease.

## Materials and Methods

### iPSC lines

The iPSC lines derived from AD patients as well as control individuals without dementia used in this study have been characterized and published (Martins et al., 2018; Schröter et al., 2016a; Schröter et al., 2016b; Schröter et al., 2016c), as detailed in Table 1. AD patients were ascertained at the memory clinic of the ZNA Middelheim, Antwerpen, Belgium in the frame of a prospective study of neurodegenerative and vascular dementia in Flanders, Belgium. Ethnicity-matched healthy individuals were screened for neurological or psychiatric antecedents, neurological complaints, and organic disease involving the central nervous system. Ascertainment and *TREM2* p.R47H genotyping are described in detail in (Cuyvers et al., 2014). iPSCs were maintained on Matrigel-coated (Corning) plates in StemMACs culture medium (Miltenyi Biotec). The medium was changed every day and the cells were passaged every 5-6 days using PBS without Calcium and Magnesium (Life Technologies).

### Neural differentiation of the iPSC lines

For the induction of GABAergic interneurons, iPSCs were differentiated using an embryoid body-based protocol (Liu et al., 2013) with modifications. On day 1, the iPS cells were harvested and re-cultivated in suspension in neural induction medium (NIM) (DMEM/F-12 (Thermo Scientific), 1% NEAA (Lonza), 1% N2 supplement (Thermo Scientific), 2 µg/ml of Heparin (Sigma-Aldrich) and 1% P/S) supplemented with 1 µM purmorphamine (Tocris Bioscience), a SHH agonist. At day 5 the formed aggregates, called embryoid bodies (EBs), were harvested and re-plated as adherent cells in the same medium and the same concentration of purmorphamine. From day 10 to 18, primitive neuroepithelia structures were formed and neural rosettes were selected with STEMDiff Neural Rosette Selection reagent (Stem Cell Technologies) and re-culture in suspension in NIM plus B27 supplement (Thermo Scientific) (without retinoic acid) and 20 ng/mL of EGF and FGF2 (both PrepoTech). After 10 days the cells maintained as aggregates (neurospheres) were dissociated into single cells with accutase (Invitrogen) and re-plated on Matrigel (Corning) for the final differentiation in neural differentiating medium (NDM) (Neurobasal 1% NEAA, 1% N2 supplement and 1% P/S) supplemented with 1µM of cAMP (Sigma-Aldrich) and 10 ng/mL of BDNF, GDNF and IGF-1 (all Immuno Tools). The iPSC-derived neurons were cultivated for approximately 80 days.

### Cryosection of Neurospheres

Neurospheres were fixed in 4% paraformaldehyde (PFA) for 30 min at room temperature, washed with PBS and cryoprotected in 30% sucrose in PBS overnight at 4°C. Subsequently, these neurospheres were transferred into embedding medium (Tissue-Tek OCT Compound 4583, Sakura Finetek), snap-frozen on dry ice and stored at -80°C. Neurospheres were cut into 10 µm thin slides using Cryostat Leica CM3050 S.

### Immunofluorescence stainings

Cells were fixed with 4% paraformaldehyde for 15 min at room temperature (RT). Neurosphere slides were thawed, dried and rehydrated in PBS. Fixed cells and neurosphere slides were permeabilized with 0.2% Triton X-100 for 10 min and blocked with 3%BSA in PBS for 1h. Samples were then incubated with the following primary antibodies overnight at 4°C: mouse anti-PAX6 (1:1000, SySy # 153011), rabbit anti-Nestin (1:400, Sigma Aldrich #N5413), mouse anti-NKX2.1 (1:1000, Millipore #MAB5460), goat anti-SOX1 (1:200, R&D # MAB3369), mouse anti-FOXG1 (1:1000, Biozol # LS-C197226), mouse anti-βIII-tubulin (1:200, Cell Signaling #TU-20), rabbit anti-MAP2 (1:1000, SySy #188002), ginny pig anti-GFAP (1:500, SySy #173004), guinea pig anti-Synapsin 1 (1:500, SySy #106004), mouse anti-SMI-3 (1:2000, Covance #SMI-312R) and rabbit anti-GABA (1:1000, Sigma Aldrich #A2052). After washing with PBS, cells were then incubated with the appropriate secondary antibody conjugated with Alexa-488, Alexa-555 or Alexa-647 (1:500, Life Technologies) for 1h at RT. The nuclear stain Hoechst 33258 (2ug/mL, Sigma) was added at the time of the secondary antibody incubation. Slices were mounted in ImmuMount (Thermo Scientific) and fluorescent images were obtained by a LSM 700 microscope (Carl Zeiss), and analyzed in Adobe Photoshop software (Adobe, USA).

### Immunoblotting of lysates from Aβ-S8C dimer stimulated cells

iPSC-derived neurons were differentiated for six weeks and then stimulated with 500 nM of oxidized Aß-S8C dimers (Müller-Schiffmann et al., 2016) for 72 h. Cells were then washed three times with PBS and then lysed in PBS/ 1% NP40 + complete protease inhibitor cocktail (Sigma) and phosphatase inhibitor cocktail 2 (Sigma). Lysates were cleared by centrifugation at 20.000g for 10min and quantified with the *DC* Protein assay Kit (Bio-Rad). 25 µg of the lysates were then separated on NuPAGE 4-12% Bis-Tris gels (Invitrogen) and blotted to a 0.2 µm nitrocellulose membrane for 2h at 400mA. The blots were blocked in PBS containing 5% skim milk and then probed with the following primary antibodies over night at 4°C: mouse anti-total tau (HT7, 1:1000, Thermo Fisher #MN1000), mouse anti-phospho tau Ser202/Thr205 (AT8, 1:1000, Thermo Fisher #MN1020), rabbit anti-APP (CT15, 1:3500, (Sisodia et al., 1993)), rabbit anti-βactin (1:5000, Sigma #A2066), mouse anti βIII-tubulin (1:1000, Cell Signaling #TU-20). After washing the blots three times with PBS/ 0,05%Tween20 they were incubated with the appropriate secondary antibody: goat anti-mouse IRDye 680RD and 800CW as well as goat anti-rabbit IRDye 680RD and 800CW (all from LI-COR Biosciences). Following three times washing with PBS/0.05% Tween20 the fluorescent signals were quantified by applying the Odyssey infrared imaging system (LI-COR Biosciences).

### Measurement of Aß_1-40_ and Aß_1-42_ by ELISA

Aß_1-40_ and Aß_1-42_ concentrations from cleared supernatants of differentiated iPSCs were quantified by using the Amyloid beta 40/ 42 Human ELISA Kits (#KHB3441 and KHB3481; Thermo Fisher) according to the manufacturer’s recommendations. Results were normalized to the protein concentration of the cells. For this the cells were washed three times with PBS and lysed in PBS/ 1% NP40. The protein content was then measured with the *DC* Protein assay Kit (Bio-Rad).

### RNA isolation and quantitative RT-PCR

Total RNA was extracted from cell lysates using Direct-zol RNA Mini Prep kit (Zymo Research) in combination with peqGOLD TriFast (PeqLab Biotechnologie) according to the manufactureŕs protocol. 0.5 μg of purified RNA was used for first-strand cDNA synthesis using TaqMan reverse transcription reagent (Applied Biosystems). cDNA was used for subsequent PCR. Real-time quantification of genes was conducted for three independent cultures from each iPSC-derived interneuron line using the SYBR® Green RT-PCR assay (Applied Biosystems). Primer sequences are provided in Supplementary Table 5 (Primers were purchased from Eurofins Genomics). Amplification, detection of specific gene products and quantitative analysis were performed using a ‘ViiA7’ sequence detection system (Applied Biosystem). The expression levels were normalized relative to the expression of the housekeeping gene *RPS16* using the comparative Ct-method 2 ^-ΔΔCt^ and the expression levels of the GABAergic interneurons markers were compared to the expression levels of the EBs, which was set to 1.

### Generation of deep sequencing data

Deep sequencing data of cDNA from iPSC-derived neuronal cultures were generated at the Neuromics Support Facility at the VIB-University Antwerpen Center for Molecular Neurology. Sequence libraries were constructed using QuantSeq 3’ mRNA-Seq Library Prep Kit (Lexogen, Greenland, NH, USA). Sequencing was performed by Illumina NextSeq sequencing. Reads were single-end with a read length of 151. Samples from two independent experiments (n= 4 cell lines) were multiplexed onto the sequencing flow cell and the measured reads were de-multiplexed for follow-up processing. Total RNA from human adult brain, human brain clinically diagnosed with AD and human fetal brain were purchased from BioChain®.

### Analysis of deep sequencing data

The de-multiplexed fastq files were aligned against the GRCh38 genome with the HISAT2 (version 2.1.0) alignment software (Kim et al., 2015) using options for clipping the 50 bases at the 3’ end of each read. The exact HISAT2 command which was mainly derived from the parameter optimizations of Barruzzo *et al*. (Baruzzo et al., 2017) was: *hisat2 -p 7 --trim3 50 -N 1 -L 20 -i S,1,0.5 -D 25 -R 5 --mp 1,0 --sp 3,0 -x hisatindex/grch38 - U input.fastq.gz -S output.sam* The resulting BAM files were sorted by coordinates applying SAMtools software (Li et al., 2009). Reads were summarized per gene with the subread (1.6.1) featurecounts software (Liao et al., 2014) against the gencode.v22.annotation.gtf using parameter –t exon –g gene_id. Summarized reads were normalized in R using the voom normalization (Law et al., 2014) algorithm from the limma package (Smyth, 2004) filtering genes which were expressed with CPM (counts per million) > 2 in at least two samples.

### Analysis of microarray data

cDNA from iPSC-derived GABAergic interneurons from CON8 and AD-TREM2-4 untreated (CTR) and treated with Aß-S8C dimer was subjected to hybridization in duplicates on the GeneChip PrimeView Human Gene Expression Array (Affymetrix, Thermo Fisher Scientific) at the BMFZ (Biomedizinisches Forschungszentrum) core facility of the Heinrich-Heine University, Düsseldorf. Data analysis of the Affymetrix raw data was performed in the R/Bioconductor (Gentleman et al., 2004) environment using the package affy (Gautier et al., 2004). The obtained data were background-corrected and normalized by employing the Robust Multi-array Average (RMA) method from the package affy. Hierarchical clustering dendrograms and heatmaps were generated using the heatmap.2 function from the gplots package with Pearson correlation as similarity measure and color scaling per genes (Warnes et al., 2015). Expressed genes were compared in Venn diagrams employing package VennDiagram (Chen and Boutros, 2011). Gene expression was assessed with a threshold of 0.05 for the detection-*p*-value which was calculated as described in the supplementary methods in Graffmann *et al*. (Graffmann et al., 2016).

### Protein interaction network

A protein interaction network was constructed from the set of 95 genes expressed exclusively in Aβ-S8C stimulated TREM2 neurons in the Venn diagram analysis. Interactions associated with *Homo sapiens* (taxonomy id 9606) were filtered from the Biogrid database version 3.4.161 (Chatr-aryamontri et al., 2017). From this dataset interactors and additionally interactors of these interactors starting at the proteins coded by the above-mentioned set of 95 genes were extracted. The resulting complex network was reduced by searching the shortest paths between the original set via the method get.shortest.paths () from the R package igraph (Csardi and Nepusz, 2006). The protein network consisting of these shortest paths was plotted employing the R package network (Butts, 2008) marking proteins from the original set in green and inferred proteins in red.

### Gene ontology and pathway analysis

Based on the set of 95 genes expressed exclusively in Aβ-S8C stimulated TREM2 neurons in the Venn diagram analysis over-represented gene ontology terms and KEGG (Kyoto Encyclopedia of Genes and Genomes) pathways (Kanehisa et al., 2017) were determined. The hypergeometric test was used for over-representation analysis – in the version from the GOstats package (Falcon and Gentleman, 2007) for GO terms and the version from the R base package for KEGG pathways which had been downloaded from the KEGG database in March 2018. Dot plots of the most significant GO terms and KEGG pathways were done via the function ggplot() from the R package ggplot2 indicating p-values from the hypergeometric test on a red-blue color scale, number of significant genes in the dedicated pathway (G) by the size of the dots and ratios of the number of significant genes in the dedicated pathway/GO to the total number of genes in that pathway/GO on the x-axis.

### Human Cytokine array

The secretion of cytokines in AD TREM2 neuronal cultures before and after stimulation with the Aß-S8C dimer was measured employing the Proteome Profiler Human Cytokine Array kit (R&D System, United States of America). The assay was performed following the manufacturer’s instructions. Briefly, AD TREM2-2 and AD-TREM2-4 cell culture supernatants from control and 72h of Aß-S8C dimer stimulation were collected, pooled and mixed with a cocktail of biotinylated detection antibodies for further incubation in a nitrocellulose cytokine array membrane with the immobilized capture antibodies spotted in duplicates. Chemiluminescent detection of the streptavidin-HRP secondary antibody was performed and the average signal (pixel density) was determined for the pair of duplicate spots using Image J (U. S. National Institutes of Health, Bethesda, Maryland, USA). The relative change in cytokine levels was performed comparing the intensity of the spots in the Aß-S8C dimer stimulated membrane with the control membrane, which was set to 100%.

### Statistical analysis

Statistical analysis was performed with GraphPad Prism Software version 6.01 (GraphPad software). For comparisons of the mean between two groups, one-tail Student’s t-test was used performed. One-way ANOVA was used for statistical significance analysis for comparisons of the mean among 4 groups, followed by a post hoc test with the use of Tukeýs multiple comparison test. Statistical significance was assumed at p<0.05. All data are expressed as mean ± standard error of the mean (SEM).

## Acknowledgments

The authors thank the Zentrum für Informations- und Medientechnologie (ZIM) at the Heinrich-Heine University for the computational support.

## Author contributions

Conceptualization: S.M. and J.A.; Methodology: S.M, A.M., M.B.; Formal analysis: S.M., A.M., W.W. J.A.; Data curation: W.W.; Investigation: S.M, A.M., M.B; Resources: K.S., C.v.B and C.K.; Writing-original draft preparation: S.M; Writing – review and editing: J.A., W.W., A.M., K.S., C.v.B, C.K.; Supervision: J.A.

---

**Supplementary Figure 1. A.** Bright field photographs representative of the iPSC-derived neuronal cultures. **B.** Relative gene expression levels of GABAergic interneurons markers (*PV, SOM, CALB2, GAD67 and GAD65*) analyzed by qRT-PCR on control and AD neuronal differentiations shown as fold change to EBs. Data are represented as mean ± SEM from three independent experiments.

**Supplementary table 1.** Gene list, GO analysis and KEGG pathways of exclusive groups and shared genes of TREM2 and CON neuronal networks shown in Figure 2.

**Supplementary table 2.** Gene list, GO analysis and KEGG pathways of the shared genes between TREM2 and CON neuronal cultures after Aβ-S8C stimulation shown in Figure 4.

**Supplementary table 3.** Relative mRNA expression of the genes enriched in the KEGG Phagosome pathway shown in Figure 5. N.S, non-significant.

**Supplementary table 4.** Gene list, GO analysis and KEGG pathways of the exclusive expressed genes in TREM2 neuronal cultures after Aβ-S8C stimulation shown in Figure 7.

**Supplementary table 5.** Relative mRNA expression of the genes enriched in the KEGG Protein Process in Endoplasmic Reticulum pathway shown in Figure 8. N.S, non-significant.

**Supplementary table 6.** Additional information about the primer sequences used in this study.

