## Supplementary figures and images for "IPSC-derived neuronal cultures expressing the Alzheimer’s disease associated rare TREM2 R47H variant enables the construction of an Aβ-induced gene regulatory network"

### Supplementary Fig.1

#### Supplementary figure 1

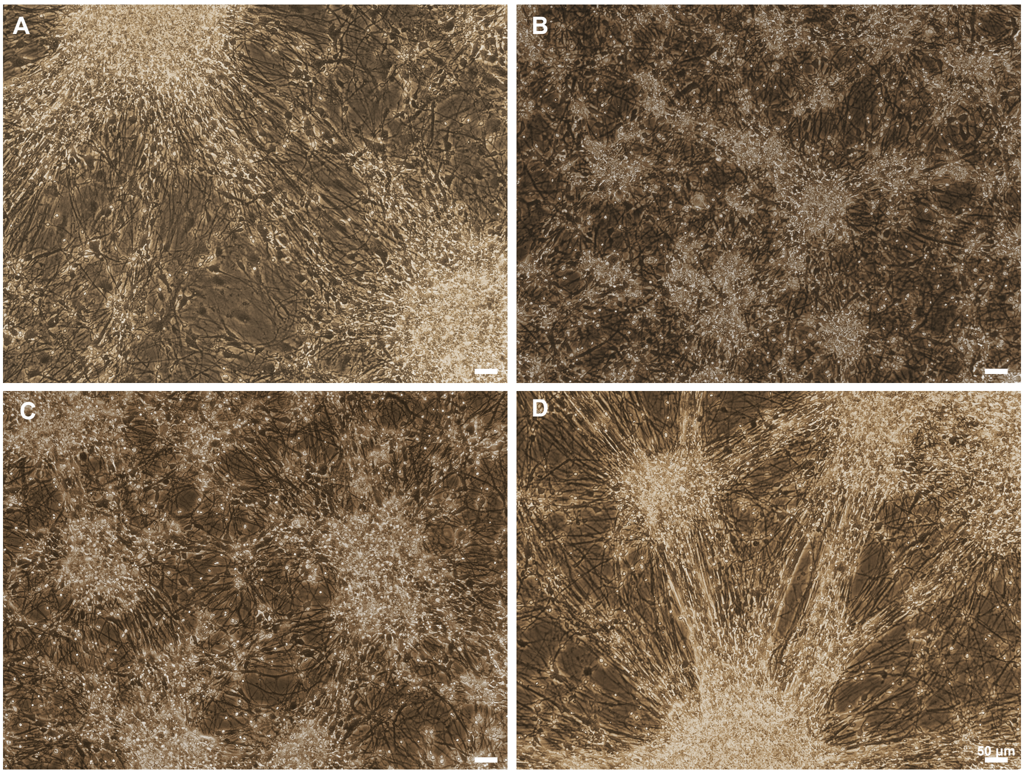


E

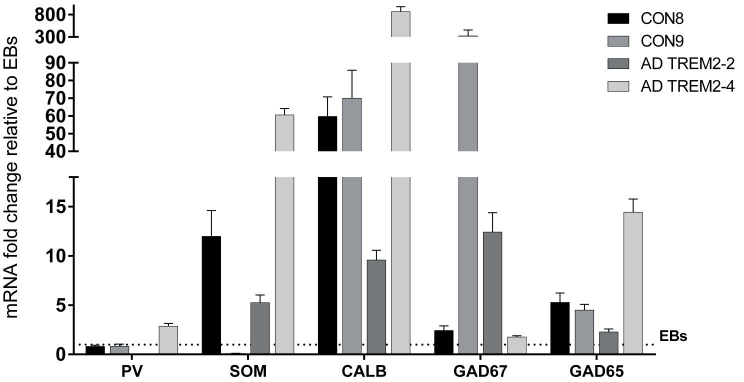
