## Supplementary Table 5 for "IPSC-derived neuronal cultures expressing the Alzheimer’s disease associated rare TREM2 R47H variant enables the construction of an Aβ-induced gene regulatory network"

| Gene name | KEGG orthology | Ratio CON8_Aβ/ CON8_CTR | Ratio TREM2_Aβ/ TREM2_CTR |
|---|---|---|---|
| XBP1 | XBP | N.S. | 3.02 |
| HERPUD1 | HERP | N.S. | 2.47 |
| DDIT3 | CHOP | 1.35 | 2.12 |
| ERP29 | PDIs | N.S. | 1.71 |
| ATF4 | ATF4 | N.S. | 1.44 |
| SSR1 | TRAP | N.S. | 1.37 |
| SEC63 | Sec62/63 | N.S. | 1.36 |
| RAD23A | RAD23 | N.S. | 1.36 |
| P4HB | PDIs | N.S. | 0.67 |
| CALR | CRT | N.S. | 0.72 |
| HSP90B1 | GPR94 | N.S. | 0.73 |
| DNAJC3 | Hsp40 | N.S. | 0.74 |
| HSPH1 | NEF | N.S. | 0.74 |
| CAPN1 | Calpain | N.S. | 0.75 |
| HSPA5 | BiP | N.S. | 0.75 |
