## Supplementary Table 6 for "IPSC-derived neuronal cultures expressing the Alzheimer’s disease associated rare TREM2 R47H variant enables the construction of an Aβ-induced gene regulatory network"

| Gene | 5'-3' Forward | 5'-3' Reverse | Template [bp] |
|---|---|---|---|
| *PV* | AAAGAGTGCGGATGATGTGAAG | ACCCCAATTTTGCCGTCCC | 186 |
| *SOM* | GCTGCTGTCTGAACCCAAC | CGTTCTCGGGGTGCCATAG | 138 |
| *CALB2* | GCTCCAGGAATACACCCAAA | CAGCTCATGCTCGTCAATGT | 208 |
| *GAD67* | AGGCAATCCTCCAAGAACC | TGAAAGTCCAGCACCTTGG | 124 |
| *GAD65* | CGCATGGTCATCTCAAACC | AGTGGAACAGCTTGGTGAGC | 114 |
| *TREM2* | TCGAGGATGCCCATGTGGAG | TTAGGAAAGACCCATCGCTGT | 146 |
| *RPS16* | GCTATCCGTCAGTCCATCTCCAA | CCTTCTTGGAAGCCTCATCCAC | 73 |
