## Supplementary Table 3 for "IPSC-derived neuronal cultures expressing the Alzheimer’s disease associated rare TREM2 R47H variant enables the construction of an Aβ-induced gene regulatory network"

| Gene name | KEGG orthology | Ratio CON8_Aβ/ CON8_CTR | Ratio TREM2_Aβ/ TREM2_CTR |
|---|---|---|---|
| HLA-DMA | MHCII | 1.33 | N.S. |
| HLA-DMB | MHCII | 1.58 | N.S. |
| HLA-DOA | MHCII | 1.37 | N.S. |
| HLA-DPB1 | MHCII | 1.38 | N.S. |
| HLA-DQA1 | MHCII | 1.46 | N.S. |
| HLA-DQB1 | MHCII | 1.57 | N.S. |
| HLA-DRB1 | MHCII | 1.34 | N.S. |
| HLA-F | MHCI | 1.60 | N.S. |
| TUBB4A | TUBA | N.S. | 0.73 |
| TUBB4B | TUBB | N.S. | 0.73 |
| DYNC1H1 | Dynein | N.S. | 0.70 |
| LAMP2 | LAMP | N.S. | 0.74 |
| ATP6V1A | vATPase | N.S. | 0.70 |
| ACTB | F-actin | N.S. | 0.69 |
| THBS1 | TSP | N.S. | 0.71 |
| CALR | CALR | N.S. | 0.72 |
| TUBA1C | TUBA | N.S. | 0.67 |
